# Parallel adaptation to geothermally-warmed habitats due to common structural variation and functional developmental pathways

**DOI:** 10.1101/2025.02.17.638600

**Authors:** Matthew K. Brachmann, Ana P. B. Costa, Shaun Robertson, Kate Donoghue, Natalie Pilakouta, Mark Whitehead, Xuan Liu, Bjarni Kristjansson, Skuli Skulason, Colin Selman, Kevin Parsons

## Abstract

Climate change is causing rapid increases in temperature which drives genomic changes tied to adaptation. However, predicting the outcomes of climate change presents challenges as the anticipated conditions have yet to be experienced by natural populations. Modelling and lab experiments suggest that natural populations will experience shifts in life history, physiology, phenology, and ecology, but the underlying genomic mechanisms involved are unknown. However, some contemporary natural populations experience habitat warming through geothermal activity and can provide valuable insights into evolutionary responses. Geothermally warmed habitats should impose strong selection on ectotherms compared to ambient habitats as they increase metabolic demands, alter developmental processes, and offer novel ecological conditions. We leveraged Icelandic threespine sticklebacks (Gasterosteus aculeatus) from populations that have adaptively diverged along a geothermal/ambient habitat axis. We obtained 173,485 single nucleotide polymorphisms (SNPs) across four independent instances of population divergence using whole genome sequencing. While the majority of genomic differentiation between geothermal/ambient ecotypes was non-parallel, the MAPK signalling pathway appeared across all ecotype pairs. We also identified a putative inversion located on chromosome XXI which appears to drive parallel genomic differentiation between geothermal and ambient ecotypes. Candidate genes within the putative inversion correspond to metabolic adaptations, including regulation of appetite and fat content. Appetite level showed strong heritable divergence between ecotypes, while the rate of weight loss during starvation and fat levels differed between ecotypes. Overall, both polygenic adaptation and parallel structural variation appeared to be key genomic mechanisms for adaptation to geothermally warmed environments. While allelic divergence was largely unique across populations, it resulted in similar functional phenotypic outcomes. Thus, structural and allelic variation both operate to facilitate adaptation to warming environments. Therefore, while management from a genomic perspective will play a role in mitigating the effects of climate change, this study suggests that consideration of functional molecular pathways will be key to conservation but with precise changes being difficult to predict due to the highly polygenic nature of thermal adaptation.

## Introduction

Increasing global temperatures represent the most pervasive threat to biodiversity (Woodward et al., 2010; Bellard et al., 2012; Haase et al., 2023). Ectotherms are predicted to be the most impacted by warming climatic conditions as the environment primarily regulates their body temperature (Diffenbaugh & Field, 2013; Kingsolver et al., 2013; Fontaine & Kohl, 2023). This will be especially challenging for populations that are limited in their ability to disperse to new habitats, as they will either be forced to adapt in situ or go extinct (Chirgwin et al., 2015). Therefore, the capacity of organisms to adapt to increased temperatures, and the mechanisms behind this, will be crucial for predicting and mitigating biodiversity loss (Franks & Hoffmann, 2012; Lancaster et al., 2022; Bernatchez et al., 2024). Thus, gaining insights into the genomic mechanisms underpinning thermal adaptation will play a critical role in mitigating climate change.

Rapid changes in climatic conditions are expected to drive adaptation through genomic mechanisms (Lancaster et al., 2022). Many studies attempting to understand the genomic basis of climate change adaptation focus on changes in allele frequencies (Franks & Hoffmann, 2012). For example, Des Roches et al (2020) showed that climate change is driving changes in allele frequencies of the EDA gene over the last century in threespine stickleback (Gasterosteus aculeatus) across a latitudinal gradient. However, while specific genes are likely to be involved, temperature has broad effects on development, physiology, and in turn gene expression and function, making it likely that thermal adaptations will be polygenic and involve complex phenotypes (Capblancq et al., 2020; Oomen et al., 2020). Across multiple populations such polygenic traits will likely emerge though unique allelic variation and be impacted by genetic background (i.e. population specific epistasis), making generalizations about the genomic basis of climate change adaptation challenging. Indeed, allelic changes are often non-parallel across populations diverging along replicate habitat gradients (Bolnick et al., 2018). Even systems with recently shared demographic histories across populations demonstrate a high degree of non-parallelism at the genomic level (Brachmann et al., 2022). Importantly, natural selection will be blind to the specific loci involved with adaptation, with the ability to only favour functional outcomes that improve fitness (Mayr, 1997).

Beyond allele frequency changes there is growing interest in other forms of genomic variation (Oomen et al., 2020; Lancaster et al., 2022; Bernatchez et al., 2024). For example, the presence of structural variants, such as inversions, can allow organisms to adapt to changing environmental conditions (Layton & Bradbury, 2022). Adaptation based on inversions is predicted to occur rapidly as they should limit the impact of gene flow by restricting recombination thereby stabilizing links among combinations of adaptative alleles within a given haplotype block (Wellenreuther & Bernatchez, 2018; Westram et al., 2022). Indeed, large inversions can drive adaptation to contemporary environmental conditions and allow for the formation of different ecotypes (Lucek et al., 2019; Todesco et al., 2020; Hämälä et al., 2021). Acting as a ‘supergene’ they can have a large effect on complex phenotypes (Mérot et al., 2021). Given the predicted speed of climate change, structural variants could promote rapid adaptation to changing thermal conditions although their role in thermal adaptation has, to our knowledge, not been demonstrated.

Predicting the evolutionary outcomes of climate change is an important challenge for genomics (Franks & Hoffmann, 2012). However, the extreme conditions predicted by climate change have not yet been experienced by wild populations. This has led researchers to compare populations from different latitudinal and altitudinal gradients (De Frenne et al., 2013; Verheyen et al., 2019). While valuable, such systems can impose confounding effects that influence evolution (Martin & MCKay, 2004; Pilakouta et al., 2020a). Conversely, lab experiments tend to involve responses that occur within a single generation rather than evolutionary change (Campbell et al., 2021). However, some extant populations can experience warming through geothermal activity and provide direct insights into responses to warming in natural systems. Geothermally warmed habitats should impose relatively strong selection on ectotherms as they are more metabolically demanding than ambient habitats, change developmental rates and processes, and offer novel ecological conditions (Kordas et al., 2011; Pilakouta et al., 2020).

Here, we leverage populations of Icelandic threespine stickleback undergoing adaptation in response to thermal habitat variation imposed by geothermal warming (Smith et al., 2024). These divergent populations exist in very close proximity, including cases of sympatry, and have repeatedly diverged between geothermal and ambient habitats throughout Iceland. This includes divergence in morphological (Pilakouta, Humble, et al., 2023), metabolic (Pilakouta et al., 2020), and behavioural traits (Pilakouta, Killen, et al., 2023; Pilakouta, O’Donnell, et al., 2023). However, the genomic basis for such adaptive divergence across thermal habitats has not been investigated.

We assessed the genomic basis for thermal habitat divergence in these populations. First, we tested for the degree of genomic differentiation between geothermal and ambient ecotypes among populations across Iceland to gain a phylogeographic understanding of divergence. Second, using our understanding of wider geographic relationships we selectively targeted populations to identify specific regions of the genome associated with parallel divergence between geothermal and ambient habitats. Third, we aimed to assess the functional and adaptive significance of genes differentiating geothermal and ambient ecotypes including phenotypic and genomic approaches. We identified a wide breadth of genetic variation involved with thermal habitat divergence which included developmental and metabolically important genes. A particularly striking finding was a structural variant differentiating geothermal and ambient ecotypes in parallel across all populations. This structural variant controls multiple metabolically important genes and appears to regulate appetite, fat stores, and starvation responses in geothermal environments.

## Results and discussion

### Phylogeographic and population genetic structure

Our results highlighted that stickleback populations were clearly differentiated by geography, in line with previous studies across environmental gradients (Rennison et al., 2020). Generally, each freshwater body formed its own branch on the distance based phylogenetic tree (Figure 1A). However, four sites (Sauðárkrókur allopatric geothermal and ambient ponds (site 1a-b), Garðsvatn (ambient lake, site 2), and Áshildarholtsvatn (sympatric sites 3-4)) within the Sauðárkrókur region of northern Iceland clustered together, despite having varied sympatric and allopatric demographic histories, likely because they are in close physical proximity to one another making shared ancestry likely (Figure 1A). Despite being geographically separated, the two marine populations (sites 15-16) were genetically similar to each other indicating a panmictic marine population acting as a reservoir of standing genetic variation (Hohenlohe et al., 2010; Kirch et al., 2021). Many of the freshwater populations were likely colonized directly from the marine environment ∼10,000 years ago during the deglaciation of Iceland (Bell et al., 1994; Einarsson et al., 2004; Aguirre et al., 2022). However, we also included water bodies formed within the last 50-70 years (sites 3 and 12c) that would have most likely been colonized from other freshwater populations of stickleback (Einarsson et al., 2004). The oldest location (Grettislaug (site 6)) was the most differentiated from all other freshwater populations and may have been one of the earliest sites colonized by marine stickleback among our populations. Common patterns of genomic divergence despite large differences in population age, relationships, and demographic history would indicate robust divergent selection between geothermal and ambient habitats. Therefore, we selected sites for further investigation which represented old (>2000 ypb, Mývatn (MYV), Grettislaug (GTS)/Garðsvatn (GAR)) and young populations (∼70 ybp, Áshildarholtsvatn (ASHN), Sauðárkrókur (SKR)) as well as allopatric (GTS/ GAR, SKR and sympatric (MYV), (ASHN)) conditions.

**Figure 1.**
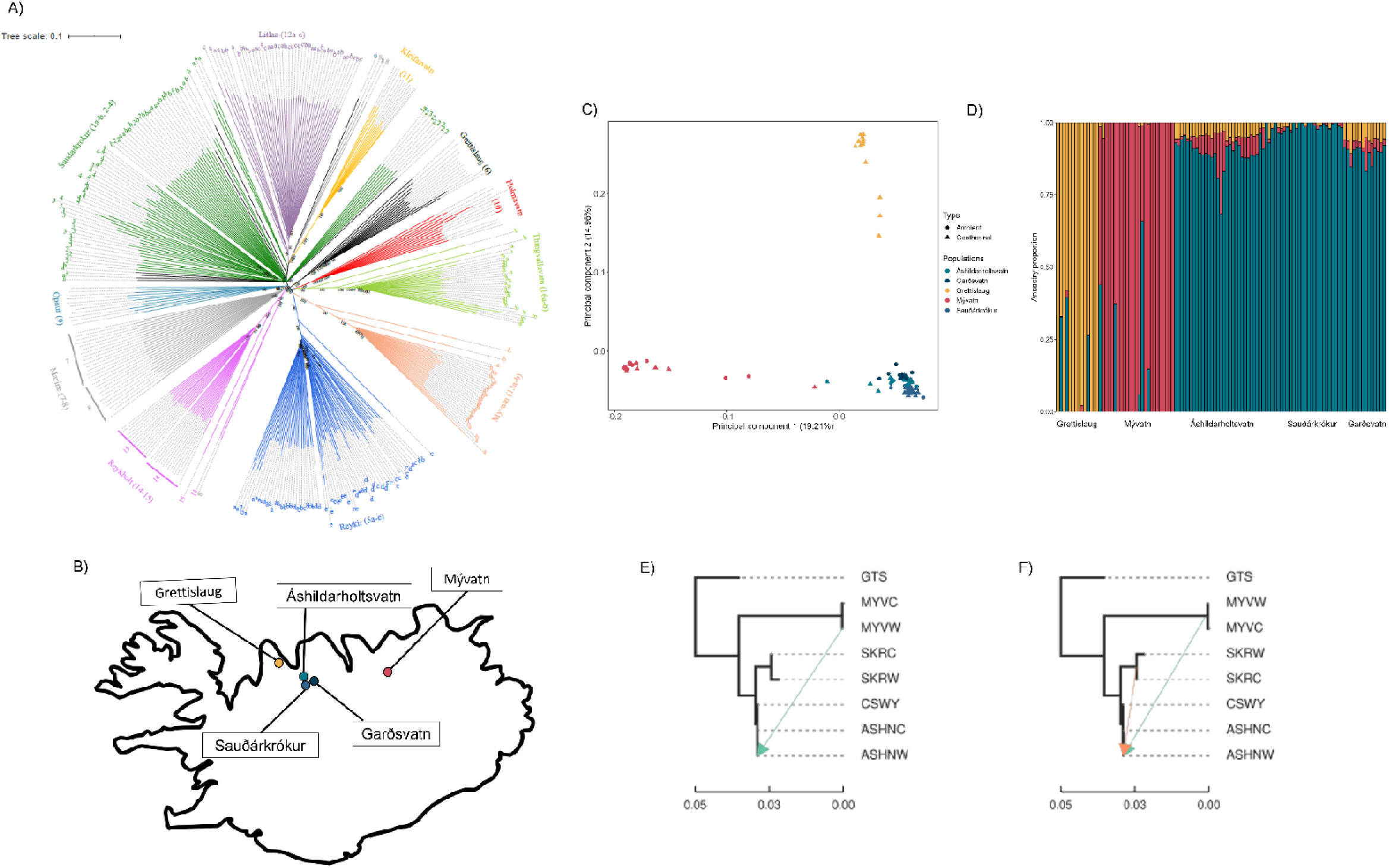
Population genetic relationships between geothermal and ambient ecotype pairs of Icelandic threespine stickleback. A) Phylogenetic relationships between 25 stickleback populations based on genotype-by-sequencing (GBS). The unrooted neighbor joining tree used a BME iterative addition method with SPR tree refinement. B) Map of Icelandic highlighting the five stickleback populations, housing four geothermal/ambient ecotype pairs, used for whole genome sequencing. C) Principal component analysis (PCA) showing the relationships between the five populations and the geothermal/ambient ecotype pairs within them. Colour codes match the map of Iceland (B). Geothermal and ambient ecotypes are shaped as triangles and dots, respectively. D) Admixture analysis showing genetic admixture across the five stickleback populations (K = 3). E) and F) Phylogenetic relationships among the five stickleback populations (Treemix) highlighting potential historical introgression between ASHN, MYV, and SKR. E) A single migration edge. F) Two migration edges.

For these populations, whole genome resequencing revealed a low degree of contemporary admixture among sites but some historical introgression was detected among some populations. We identified three distinct clusters (K = 3 corresponding to GTS, MYV, and sites in the Sauðárkrókur region) across Iceland encompassing geothermal and ambient ecotypes diverging in both sympatric and allopatric conditions (Figure 1C-D). Among these clusters there was little admixture however, there was some evidence of historical introgression between MYV and ASHN (Figure 1E) and ASHN and the SKR populations (Figure 1F). Such historical introgression events can promote shared genomic variation (alleles or structural variants) across populations and parallel genomic adaptation to contemporary environments (Kawakami et al., 2011; Lait et al., 2024).

### Local adaptation to geothermal habitats is polygenic

Genome wide genomic differentiation (Fst) between geothermal and ambient ecotypes was low across all populations but varied (MYV = -0.0022; ASHN = 0.0027; SKR = 0.012; GTS-GAR = 0.050). Allopatric sites had greater genomic differentiation between ecotypes than sympatric sites likely due to a lack of gene flow and promotion of genetic drift (Yeaman & Whitlock, 2011; Feder et al., 2012, 2013, 2014). While reduced gene flow does aid in promoting increased genomic differentiation, differences in thermal habitat appeared to be the driver of genomic differentiation between ecotypes across all population pairs.

Indeed, despite low overall genomic differentiation between ecotypes across population pairs, we detected extensive per-locus genomic differentiation between thermal ecotypes regardless of population age and demographic history. Genomic differentiation between thermal ecotypes is highly polygenic involving hundreds of outlier loci (ASHN = 281; MYV = 267; SKR = 292; GTS-GAR = 351) across multiple chromosomes (Figure 2A-D). This supports the idea that polygenic adaptation for complex adaptive phenotypes is more common than previously thought (Yeaman, 2022). We also detected regional Fst peaks differentiating geothermal and ambient ecotypes across the genome (Figure 2A-D). However, among these there was a low degree of parallelism in Fst outlier loci across geothermal and ambient ecotype pairs (Table 1; below diagonal). Notably, the youngest (SKR, ∼70 generations) and the oldest population pairs (GTS-GAR, ∼ 4000 generations) shared the greatest overlap in Fst outliers (3 shared Fst outliers) indicating that variation in age did not provide a strong influence on thermal habitat divergence. Consistent with low parallelism in outlier loci, we also detected low parallelism across ecotype pairs at the gene level (Table 1; above diagonal). Genomic parallelism in cases of divergent parallel phenotypes across populations has been shown to increase at functional levels of the genome (Brachmann et al., 2022; Jacobs et al., 2020). This contrasts with previous findings that stickleback ecotypes diverging across other environmental gradients have a high degree of parallelism at a single locus (Rennison et al., 2019; Des Roches et al., 2020) as well as multiple loci (Stuart et al., 2017; Garcia-Elfring et al., 2021). The genomic architecture of polygenic adaptation to thermal habitats is largely non-parallel across independent ecotype pairs. However, low genomic parallelism does not preclude the possibility that selection favours similar biological function. Indeed, genetic redundancy can lead to highly parallel molecular pathway function and ultimately parallel phenotypes (Barghi et al., 2019, 2020; Láruson et al., 2020).

**Figure 2.**
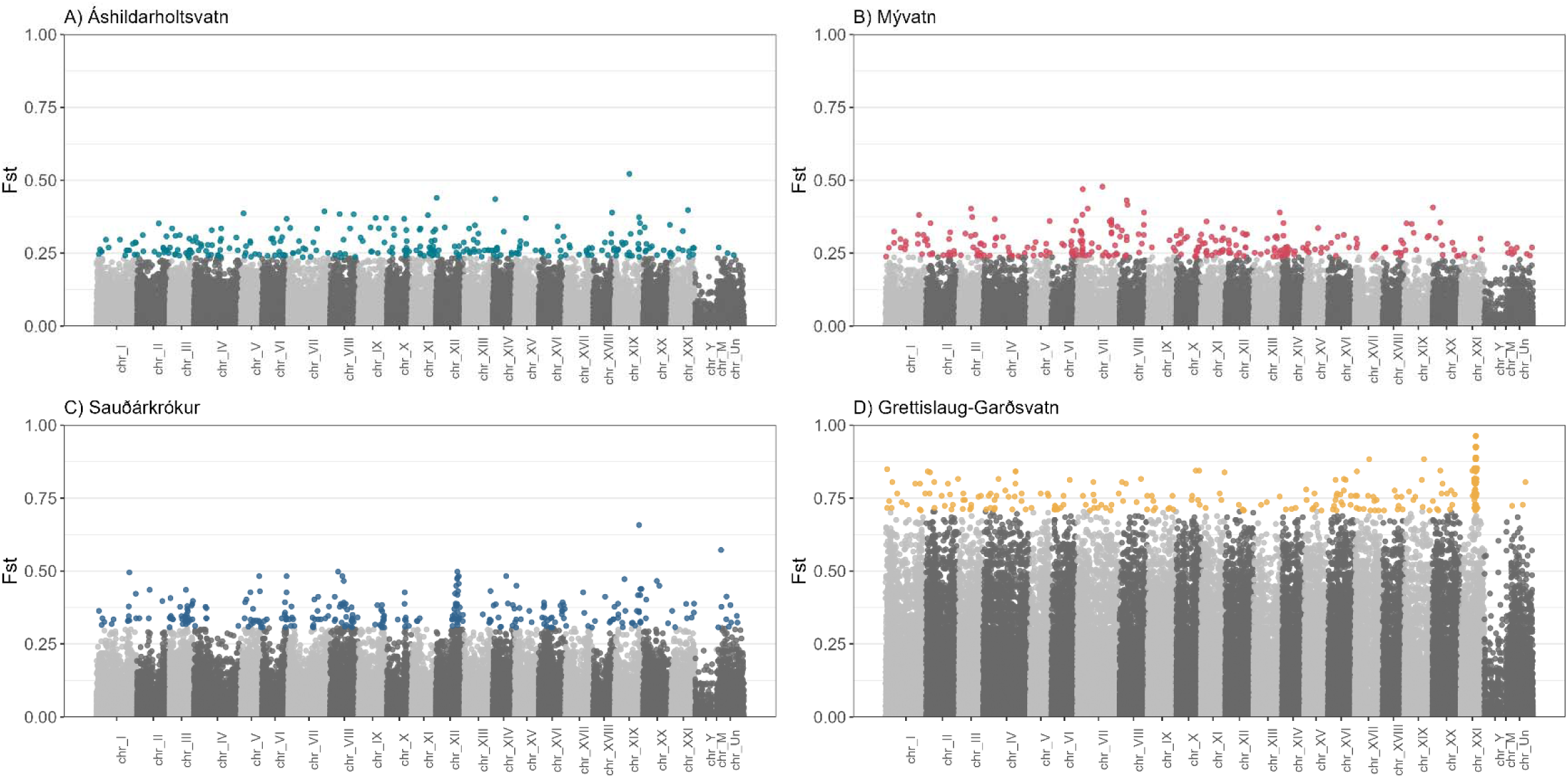
Per locus population differentiation (Fst) across the stickleback genome for four independent geothermal/ambient ecotype pairs. Neutral loci are coloured as shades of grey while Fst outlier loci are coloured according to the ecotype pair. Fst outlier loci were in the top 0.5% of the Fst distribution.

**Table 1.**
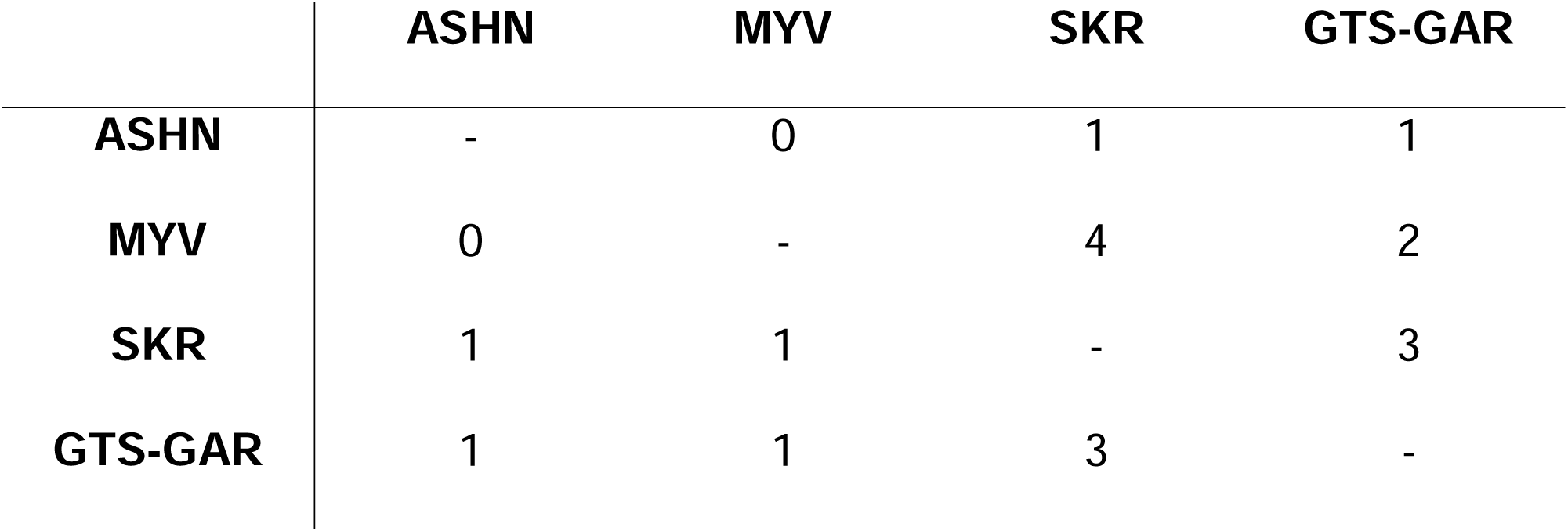
Overlap between Fst outlier loci and genes associated with Fst outlier loci across geothermal and ambient ecotype pairs of Icelandic threespine stickleback. The number of overlapping Fst outlier loci are shown below the diagonal while the number of overlapping genes associated with Fst outlier loci are show above the diagonal.

|  | ASHN | MYV | SKR | GTS-GAR |
| --- | --- | --- | --- | --- |
| ASHN | - | 0 | 1 | 1 |
| MYV | 0 | - | 4 | 2 |
| SKR | 1 | 1 | - | 3 |
| GTS-GAR | 1 | 1 | 3 | - |

Despite low genomic parallelism, we found evidence of parallel functional divergence of molecular pathways. There was a high degree of congruence in enriched gene ontology (GO) terms, biological processes, and gene networks among the geothermal/ambient ecotype pairs. The enriched terms from our GO analyses for each ecotype pair showed strong commonalities, although also some differences (Figure 3 A-D). Specifically, three of the four ecotype pairs had enriched GO terms for nerve and muscle cell development, differentiation, and proliferation. Increasing temperatures in freshwater habitats have been known to cause divergence in muscle fiber development and type of muscle produced (i.e. red vs. white muscle) which in turn affects swimming performance and collagen metabolism in contrasting thermal regimes (Rome, 1990; Johnson & Johnston, 1991; Lin et al., 2022). The nervous system is also highly sensitive to differences in thermal conditions, as increased temperatures during development increases neuronal activity and growth (Beltrán et al., 2021). While further research is required, our findings help provide new insights into potential mechanisms for the reduced sociability observed in these stickleback populations (Pilakouta, O’Donnell, et al., 2023). For the oldest (∼4000 ybp) ecotype pair (GTS-GAR), genes were mainly associated with metabolism and metabolic regulation. This matched with our previous findings (Pilakouta et al., 2020) of metabolic divergence within the ecotype pair, but also across additional pairs (MYV and ASHN). Thus, while GO analysis did not provide an exact match across population pairs these findings from the oldest pair highlight functional changes that could be relevant to all ecotype pairs.

**Figure 3.**
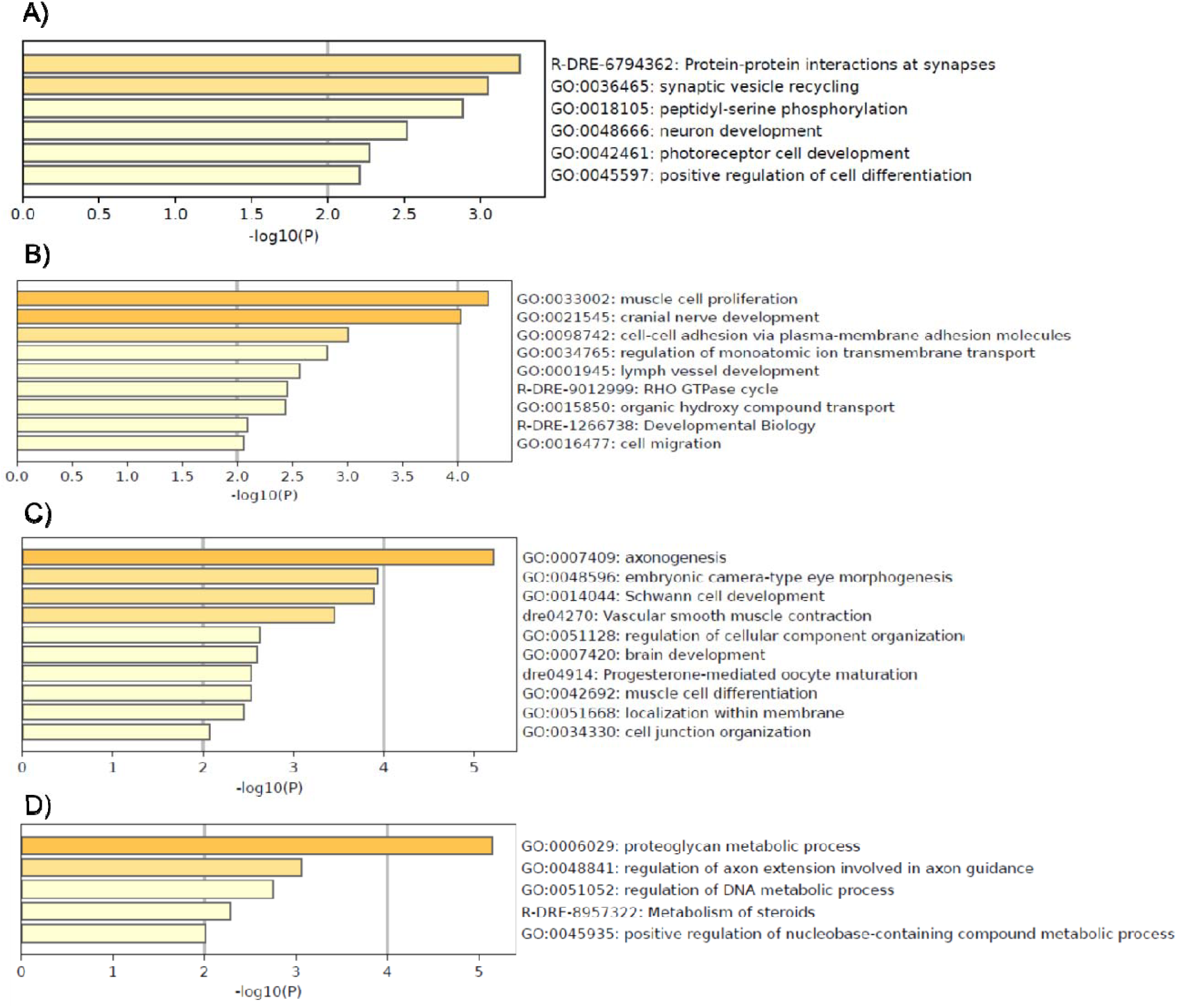
Gene ontology enriched terms from differentiated genes between geothermal and ambient ecotypes of Icelandic threespine stickleback. A) Áshildarholtsvatn geothermal and ambient ecotypes. B) Mývatn geothermal and ambient ecotypes. C) Sauðárkrókur geothermal and ambient ecotypes. D) Geothermal ecotypes from Grettislaug and ambient ecotype from Garðsvatn.

The low degree of genomic parallelism coupled with largely common functions in GO terms indicates a complex genomic basis for thermal phenotypes. This is supported by our findings of highly polygenic genome-wide differentiation as the average number of genes differentiating geothermal and ambient ecotypes was 113 (ASHN = 108; MYV = 103; SKR = 124; GTS-GAR = 120). Between 34.3 to 45.5% of outlier loci were located within non-coding regions (ASHN = 38.4%; MYV = 37.3%; SKR = 42.5%; GTS-GAR = 34.2%) while 17.6% to 24.2% were within protein coding regions of exons (ASHN = 17.6%; MYV = 16.5%; SKR= 19.3%; GTS-GAR = 24.2%). A low proportion of outlier loci were located within promoter regions (5̕ untranslated region) (ASHN = 0.93%; MYV = 0.97%; SKR = 0.81%; GTS-GAR = 1.7%) and 3̕ untranslated regions (ASHN = 0.93%; MYV = 1.94%; SKR = 0.81%; GTS-GAR = 0.83%). Much of the genomic differentiation between ecotypes appears to be driven by non-coding and potentially regulatory regions of the genome, a finding in line with previous comparisons between marine and freshwater stickleback where only 17% of divergent SNPs were within coding regions of genes (Jones et al., 2012). Changes in regulatory regions in genes associated with adaptive divergence may be more prevalent than previously thought and may allow for the divergence of complex genomic networks and cellular pathways (Jeukens & Bernatchez, 2012; Okude et al., 2024). The rates of evolutionary change are typically increased in either regulatory or coding regions but rarely both simultaneously (De-Kayne et al., 2024). Given the large scale of polygenic changes to regulatory functions of genes, as well as the non-parallel nature of adaptive divergence, we suggest that predictive allelic responses to increased warming could be intractable (Schlötterer, 2023). Thus, gene-targeted conservation strategies will likely fall short of conserving species under changing climatic conditions (Kardos & Shafer, 2018), and conservation genetic/genomic strategies should consider polygenic models of adaptation and target functional molecular pathways capable of producing adaptive phenotypes.

Our gene network analyses for each geothermal and ambient ecotype pair identified a range of putative candidate pathways (Table 2; Supplementary figure 2). Of particular interest were genes associated with the MAPK signalling pathway which appeared prominently across ecotype pairs and included specific MAPKgenes (MAP2K5 in ASHN; MAPK1 in MYV and SKR; MAPK13 in SKR). These genes have all been implicated in cell proliferation and neural crest cell development (Dinsmore & Soriano, 2018) and in turn craniofacial development (W. Zhang & Liu, 2002; Greenblatt et al., 2013). This matches with our previous findings of parallel morphological divergence in the craniofacial apparatus across ecotype pairs (Pilakouta, Humble, et al., 2023). Such morphological divergence is both heritable and adaptive across thermal habitats (Pilakouta, Humble, et al., 2023; Smith et al., 2024). While neural crest cells have been widely reported as a mechanism underlying adaptive variation (Brandon et al., 2023), it is notable that they are also sensitive to temperature variation with activated ion channels conferring maternal fever associated birth defects (Canales Coutiño & Mayor, 2022). Environmental variation in temperature can facilitate differences in the timing of neural crest cell differentiation and proliferation as well as differences in cell type differentiation (Canales Coutiño & Mayor, 2022). Thus, while more investigation is required, neural crest cells may play a large role in thermal adaptation.

**Table 2.**
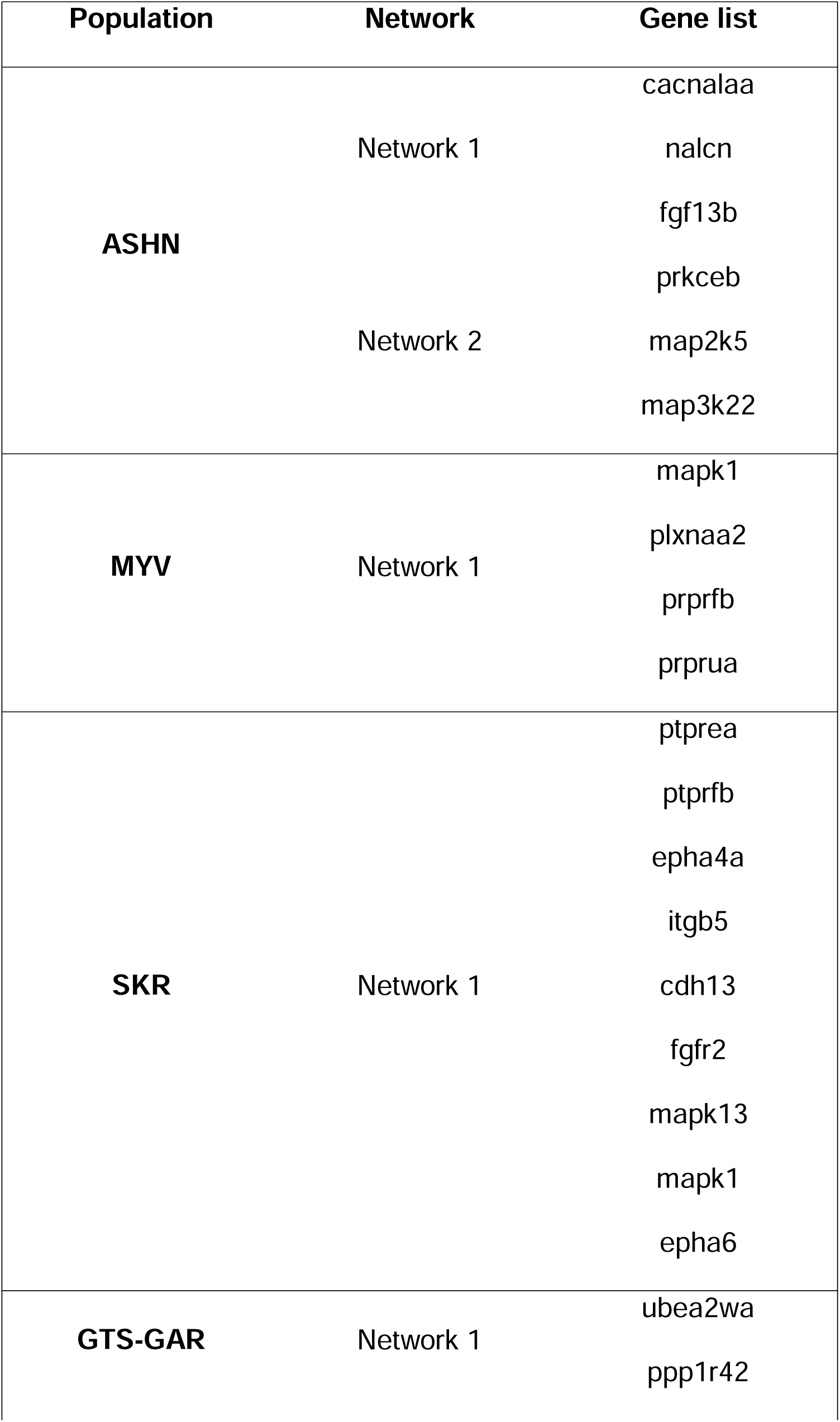
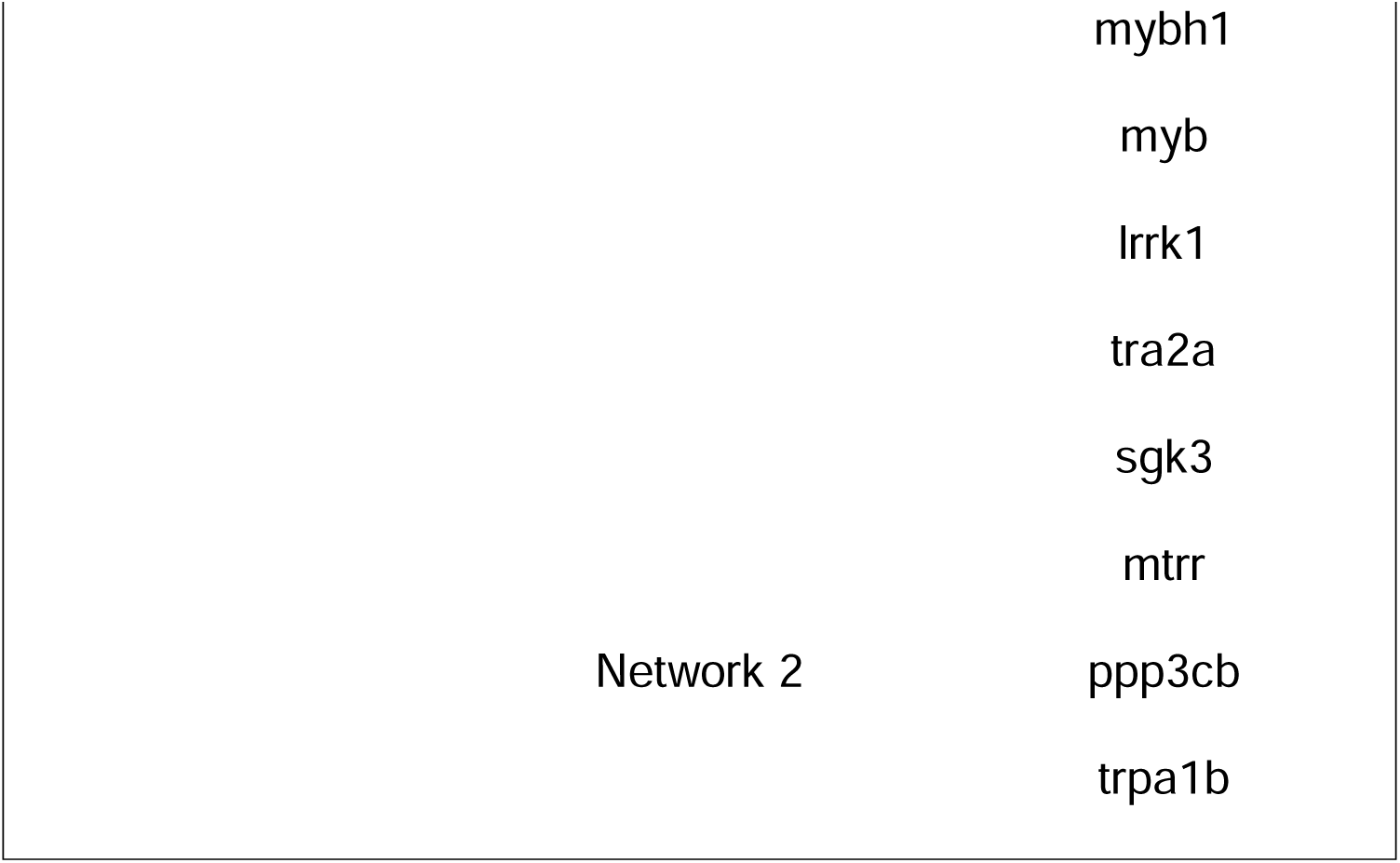
Networks of genes which have proteins that form physical interactions for each geothermal-ambient ecotype pair across five populations of Icelandic threespine stickleback. Gene lists were formed for each geothermal-ambient ecotype pair based on their physical location to Fst outlier loci.

### Parallel divergence between geothermal and ambient habitats relies on structural variation

Despite low levels of genomic differentiation (interspersed with outliers) we found strong evidence of parallel differentiation in a localized but multigene region. Specifically, a 1.6 megabase (Mb) region on chromosome XXI showed both elevated and common allele frequency divergence between thermal ecotypes (Figure 4C; Supplementary Figure 5). Within this 1.6Mb region there was a distinct Fst peak differentiating thermal ecotypes (Figure 4B). Overall, divergence was relatively high with 46.2% of all loci on chromosome XXI being Fst outliers, and with 64% of these loci being located within the 1.6Mb region (Figure 5A). This divergent region of chromosome XXI showed elevated levels of the XP-EHH statistic (Figure 5B) suggesting strong selection associated with the geothermal environment. Lastly, this region of chromosome XXI was identified as an MDA outlier (Figure 5C) indicating structural variation such as an inversion.

**Figure 4.**
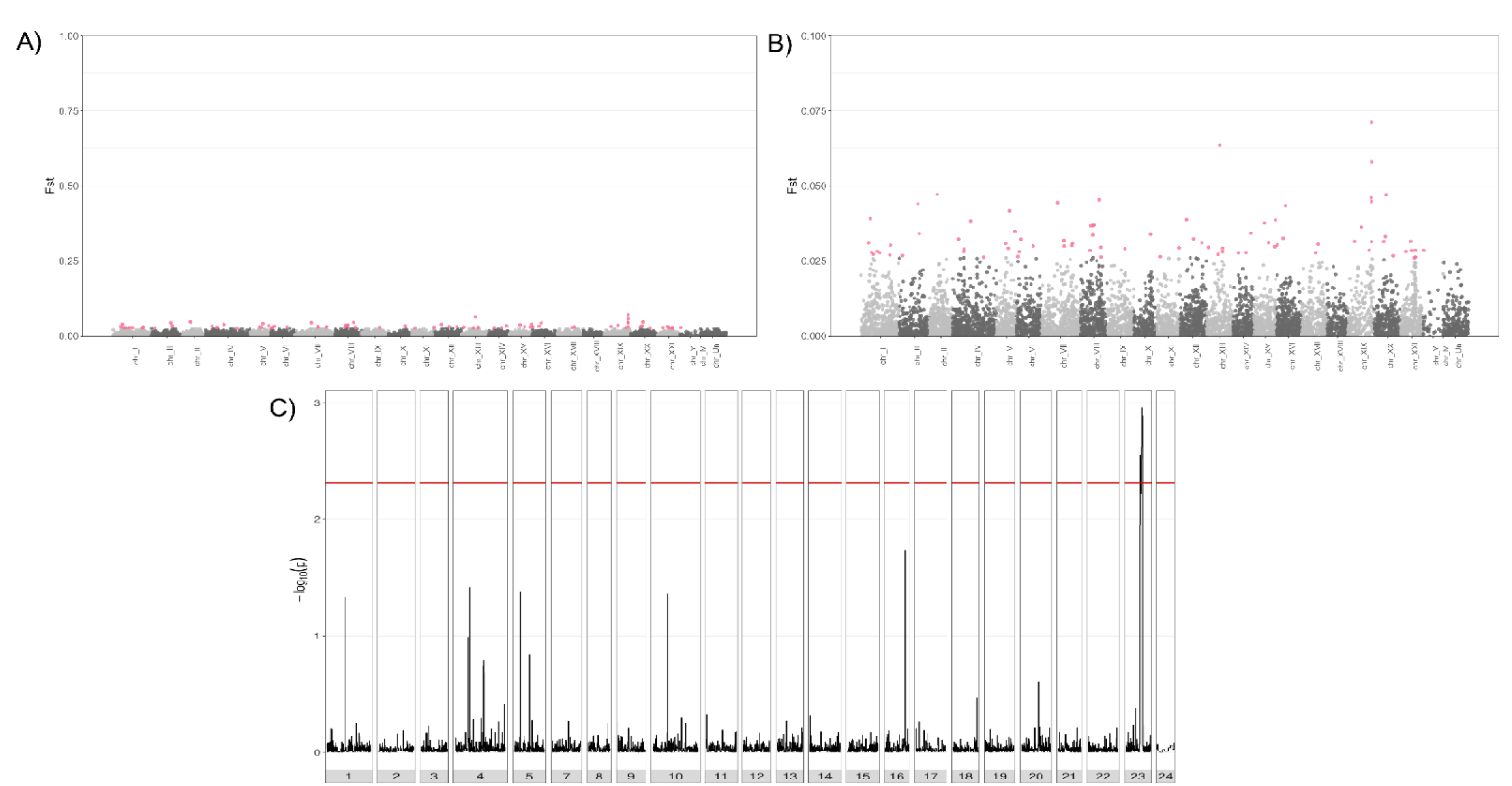
Assessment of parallel differentiation of geothermal and ambient ecotype pairs of Icelandic stickleback. A) Genome-wide differentiation (Fst) between pooled geothermal and pooled ambient ecotype pairs based on 25 kilobase sliding windows. B) Genome-wide differentiation (Fst) between pooled geothermal and ambient ecotype pairs based on per locus Fst. Outlier loci are coloured while neutral loci are shades of grey. C) Parallel differentiation of allele frequencies between geothermal and ambient ecotype pairs(AF-vapeR). The red line indicates significance based on a null distribution of allele frequencies. Significant peaks of parallel differentiation in allele frequencies are located on chromosome XXI.

**Figure 5.**
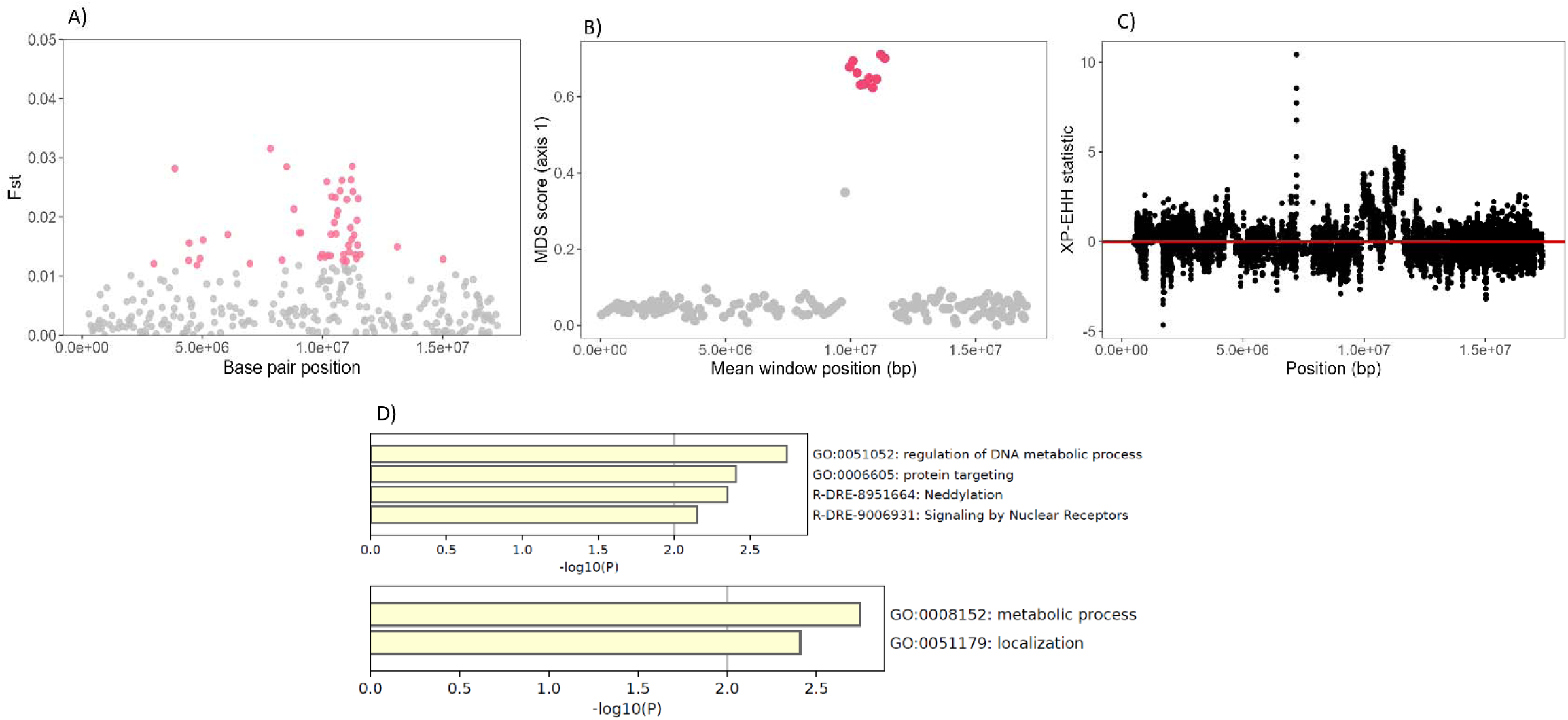
Assessment of the type of genomic variation represented by the parallel region on chromosome XXI differentiating geothermal and ambient ecotype pairs. A) Per locus differentiation between all geothermal and ambient ecotype pairs. Outlier loci are coloured while neutral loci are represented in grey. B) Differentiation of population structure between geothermal and ambient ecotype pairs along chromosome XXI in 50 SNP windows (lostruct). Significant deviations in population structure are coloured while neutral windows are represented in grey. Deviations in population structure using local PCA have been previously indicative of structural variants. C) Putative selective sweeps between geothermal and ambient ecotype pairs along chromosome XXI. The red line indicates neutrality, positive deviations indicate selective sweeps in geothermal ecotypes, negative deviations indicate selective sweeps in ambient ecotypes. D) Identification of biological functions for the genes within the region of interest on chromosome XXI using gene ontology (GO). The top panel shows enriched terms while the bottom panel shows the biological function for the 55 genes within the region on chromosome XXI.

Structural variants have been implicated in promoting adaptive divergence among ecotypes as large regions of co-adapted genes can be inherited as a single unit (Mérot et al., 2020; Oomen et al., 2020) and thus effectively act as a single locus that is resistant to the effects of recombination (Wellenreuther & Bernatchez, 2018; Lucek et al., 2019; Wellenreuther et al., 2019). The structural variant on chromosome XXI has previously been inferred as an inversion associated with the transition from marine to freshwater environments in threespine stickleback (Jones et al., 2012; Dean et al., 2024). As all stickleback originated from freshwater environments, we instead suggest this inversion is related to transitions from cold to warm thermal habitats across the species. Marine and freshwater environments differ not only in salinity but also temperature profiles as freshwater habitats are generally more variable and achieve warmer temperatures than marine habitats (Hölker et al., 2004; Comte & Olden, 2017). Wild marine and freshwater stickleback differ in their ability to handle extreme temperatures as marine individuals have impaired cold tolerance relative to freshwater individuals (Barrett et al., 2010). However, while cold tolerance can evolve rapidly (within three generations) heat tolerance does not, suggesting a more complex genetic basis for adaptation to warmer temperatures. Therefore, understanding the effects of this inversion on thermal habitat divergence could provide more insight into the function of inversions, including the speed at which thermal adaptation can occur as it is implicated in both young and old population pairs.

The putative inversion contained genes relevant to thermal adaptation, including metabolism, morphology, neddylation, and epigenetic regulation (Figure 5D). Gene ontology analysis on the 55 genes in this region matches with our previous findings implicating morphology and metabolism as divergent traits in these systems (Pilakouta et al., 2020; Pilakouta et al., 2023). More broadly for threespine stickleback, previous independent studies have identified 23 morphological QTL within this region that are associated with feeding, body shape, defence, and sensory systems (Wark et al., 2012; Erickson et al., 2016; Liu et al., 2016; Peichel & Marques, 2017). Geothermal environments are stressful and promote strong selection for adaptations which increase performance under stress (reference). Thermal stress can induce DNA damage and cellular apoptosis (Cheng et al., 2018; Burraco et al., 2020). Neddylation, among other processes, has been shown to minimize the effects of direct or indirect DNA damage (Brown & Jackson, 2015; S. Zhang et al., 2024). Neddylation could allow for repair of deleterious genomic mutations induced by living in geothermal environments. Geothermal environments have also been shown to induce elevated metabolic rates in ectothermic organisms (Kingsolver et al., 2013). For metabolism in particular, the geothermal habitats should cause long periods of low resource availability during winter, but alongside elevated metabolic demands due to higher temperatures. The seasonal metabolic demands faced within geothermal habitats is similar to the seasonal metabolic demands shown to influence divergence in Mexican cave fish (Astyanax mexicanus) (Cobham & Rohner, 2024). Mexican cave fish have shown remarkable adaptation to seasonal resource limitation through changes in a suite of metabolic processes (Krishnan et al., 2022; Xiong et al., 2022; Olsen et al., 2023; Pozo-Morales et al., 2024). The 1.6Mb inversion located on chromosome XXI potentially enables enhanced DNA repair and metabolic adaptation to thermal stress under geothermal conditions.

### Connecting adaptive phenotypes to candidate genes

While several candidate genes for geothermal/ambient divergence exist within the putative inversion region, some were of particular interest. This included the presence of Nuclear Receptor Coactivator 2 (NCOA2) which has been linked to appetite regulation (Ramayo-Caldas et al., 2014; Lu et al., 2015; Rosen, 2016; Krishna et al., 2018) and lipogenesis (Lu et al., 2015). This occurs through a regulatory process where NCOA2 binds to Peroxisome Proliferator Activated Receptor (PPARA) as constituent parts of the pathway associated with appetite (Rosen, 2016). PPARA has previously been shown to regulate lipogenesis in Mexican Cavefish (Xiong et al., 2022) as part of an adaptive strategy to alleviate long periods of low food availability. Thus, an NCOA2/PPARA interaction could be regulating appetite levels and lipogenesis/obesity in geothermal ecotypes. While we have not directly tested this potential link, we have found strong heritable differences in appetite between ecotypes where geothermal fish exhibit hyperphagia with much high appetite levels than ambient ecotypes (ASHN = 53.9% greater; MYV = 63% greater; SKR = 35.8% greater appetite) (F-value: 27.40; p-value = <0.001, Figure 6A). Lipogenesis also seems to differ between ecotypes as geothermal fish lose mass at a slower rate than ambient ecotypes when food intake is reduced (F-value = 6.47; p-value = 0.013) (Figure 6B & C). Geothermal ecotypes also increase fat content more rapidly during the summer season relative to ambient fish (Figure 6D). NCOA2 variants and their interaction with PPARA are potential candidates regulating increased appetite (hyperphagia) and lipogenesis in the geothermal ecotypes. Hyperphagia could be adaptive in geothermal fish leading to increased lipogenesis and energy reserves for winter periods of low food availability while under high metabolic demands due to warm temperature. Hyperphagia has previously shown to be adaptive in Mexican cavefish as a means to bridge periods of resource limitation (Aspiras et al., 2015) while increased appetite has been linked to increased obesity in most vertebrates (Berthoud & Morrison, 2008).

**Figure 6.**
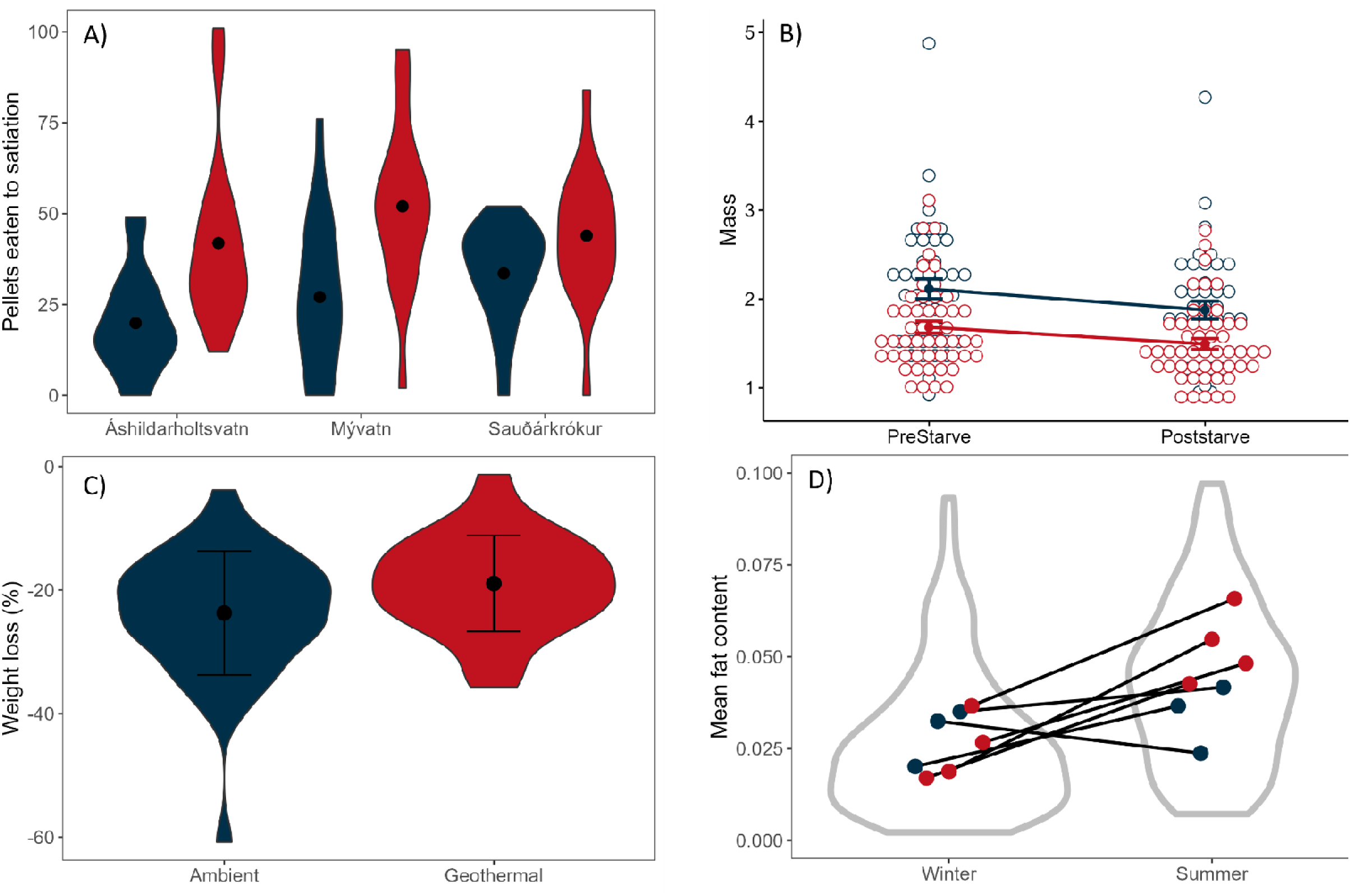
Phenotypic differences between geothermal and ambient Icelandic stickleback ecotypes. Geothermal and ambient ecotypes are shown in red and blue, respectively. A). Differences in appetite phenotypes in F2 progeny shown by the number of pellet food eaten from ASHN, MYV, and SKR. The mean number of pellets eaten is indicated by the black dot. B) Mass differences between geothermal and ambient ecotypes pre- and post-starvation for 30 days. C) Percent weight loss post-starvation for geothermal (red) and ambient (blue) ecotypes. Points represent the mean and the error bars represent standard deviation. D) Fat content of geothermal and ambient ecotypes from five Icelandic populations between summer and winter seasons. Points are represent the mean fat content for each ecotype across the two season and the violin plots show the distribution of fat content between the season across all ecotypes.

## Conclusion

Warming of natural habitats will be of increasing concern over the next decade. Here, we have provided direct evidence for a range of genomic responses to changes in thermal habitat variation in a series of natural populations of threespine stickleback. Overall, genomic differentiation between geothermal and ambient ecotypes is generally non-parallel at specific loci, but specific gene families (in this case mapk genes) provide functional similarity in molecular pathways across independent populations. However, a large putative inversion shows strong evidence of parallel divergence and involves genes relevant to a range of adaptive phenotypes. Particularly, within this region NCOA2 provides a logical candidate for divergent appetite levels and lipogenesis between geothermal and ambient ecotypes. Together these findings demonstrate that climate change adaptation is likely to involve complex genomic changes that are largely population specific. Thus, targeting certain genomic changes may fail to conserve species in warming environmental conditions, but an approach focused on conserving particular molecular pathways may be more efficient (Barghi et al., 2020). Structural variants, like inversions, may be more prevalent than previously thought in promoting adaptive divergence in this context and may help to mitigate biodiversity loss due to climate change. The complexity of such genomic changes may pose issues of tractability for conservation genomic management. Investigating the relative influence, and interactions, between structural variants and allelic variation on phenotypic adaptation may provide some resolution.

## Methods

### Study sites and sampling

To gain a phylogeographic understanding of the genetic variation occurring across populations, threespine Stickleback were collected from twenty three freshwater populations and two marine populations across Iceland (n = 365) (populations were defined based on geographic location of sites). Geothermal habitats generally had temperatures greater than twenty degrees Celsius while ambient habitats were roughly ten degrees colder within the summer months (Pilakouta et al., 2023). Freshwater populations were also classified as sympatric or allopatric populations depending on the degree of connectivity between geothermal and ambient locations. Sympatric populations allowed for continuous migration or gene flow while allopatric populations had a physical barrier separating geothermal and ambient habitats. Stickleback were sampled between May and June in 2016 and 2017 using minnow traps. Fish were humanely euthanized and fin clips taken were placed in 100% ethanol. All fish were handled following UK Home Office guidelines for animal use.

### Genotyping-by-sequencing to determine phylogenetic relationships

The genomic DNA was extracted from fin clips taken from all individuals sampled using either the Qiagen DNeasy blood and tissue kit or the Applied Biosystems MagMAX DNA Multi-sample kit used on the Kingfisher Flew automated DNA extraction machine following the manufactures protocols. Quality of the extracted DNA was assessed using a Nanodrop 8000 spectrophotometer and a 1% agarose gel while DNA concentration was measured using a Qubit BR assay in a Qubit 2.0 Fluorometer.

Genotype-by-sequencing (GBS) library preparation was conducted using the genomic DNA of all stickleback samples. GBS sequencing libraries were constructed following the protocols outlined in Elshire et al., (2011) with some modifications. A total of 100 ng of genomic DNA was digested with ApeKI and custom adapters ligated to the resulting sequence overhang. The custom adapters were designed using a dual indexing approach to control for index hopping. Samples were pooled and a double sided size selection step was performed with AMPure AP beads using a bead:sample ration of 0.7:1 followed by a 1:1 ratio for an insert size range of 100 – 400 basepairs (bp). Polymerase chain reaction (PCR) was performed in a 50 μl reaction with 25 μl NEBNNext Ultra II mastermix, 25 μM of each primer, and 20 μl of pooled library. The PCR profile was 30 seconds at 98 degrees Celsius followed by 5 cycles of 10 seconds at 98 degrees Celsius, 30 seconds at 70 degrees Celsius, 45 seconds at 65 degrees Celsius, with a final extension of 5 minutes at 65 degrees Celsius. Libraries were analysed on an Agilent Bioanalyzer High-Sensitivity Chip for quality control. Paired-end sequencing with 150 bp read length was conducted on an Illumina HiSeq 4000 sequencer at the Center for Genomic Research, University of Liverpool.

Raw reads were de-multiplexed with GBSX v1.3 (Herten et al., 2015) using the unique dual indices per sample. Any reads not containing the possible barcode combinations were removed. FastQC v0.11.7 (Andrews, 2010) was used to determine the quality of the sequence data. The reads were filtered and trimmed using Trimmomatic v0.36 (Bolger et al., 2014).

Leading and trailing bases were set to 3, the minimum phred score was set to 20, the sliding window to 4 bp, and reads less than 30 bp were removed. Reads were aligned to the Stickleback reference genome (Nath et al., 2021) using BWA-MEM v0.7.17 (Li & Durbin, 2009). The alignment bam files were sorted and indexed with Samtools v1.6 (Li et al., 2009). Samples were re-aligned around indels using GATK v4.1.3.0 (Zhou et al., 2024) with the functions RealignerTargetCreator and Indelrealigner using their default parameters. Quality of the mapped reads were checked with the functions flagstat and depth in Samtools and samples that did not pass the quality-check were removed from downstream analyses.

Biallelic single nucleotide polymorphisms (SNPs) were called using ANGSD v0.929 (Korneliussen et al., 2014). ANGSD allows for calculating genotype likelihoods for each individual which accounts for uncertainty due to low sequencing depth (Korneliussen et al., 2014).

Genotype likelihoods were estimated using the Samtools GL model and the major and minor allele frequencies were estimated with the EM algorithm (Kim et al., 2011). Only sites with significant evidence for polymorphism (P < 1×10^-6^) and a minor allele frequency (MAF ≥ 0.05) were included. Read quality filters were set using minimum mapping (minMapQ ≥ 30) and base qualities (minQ ≥ 20). Only sites present in at least 80% of the samples were kept. We excluded the sex chromosome (chromosome 19) to avoid potential sex biases in the downstream analysis.

The genetic relationships among all individuals and populations was explored with pairwise genetic distances estimates with ngsDist v1.0.8 (Vieira et al., 2016) with branch support based on bootstrapping 500 replicates with 500 SNP blocks and pairwise deletion. The phylogenetic tree was inferred with FastME v2.1.5 using a BME iterative taxon addition method with SPR tree refinement (Lefort et al., 2015). Support values were mapped using RAxML-NG v0.9.0 (Kozlov et al., 2019). The final bootstrapped tree was visualized and plotted using the online software iTOL v5 (Letunic & Bork, 2021).

### Whole genome sequencing

Using the phylogenetic tree as a reference (Figure 1A), we strategically targeted four divergent geothermal/ambient ecotype pairs for whole genome sequencing. The four geothermal/ambient ecotype pairs were subsampled based on variation in their demography (sympatry vs. allopatry) as well as their degree of phylogenetic divergence. Grettislaug (GTS) and Garðsvatn (GAR) are an evolutionary old allopatric pair of geothermal and ambient ecotypes, respectively. Mývatn (MYV) represented a pair of evolutionary old sympatric geothermal and ambient ecotypes. Áshildarholtsvatn (ASHN) represented a pair of evolutionary young (diverged within the last 70 years) sympatric geothermal and ambient ecotypes. Lastly, Sauðárkrókur (SKR) represented a pair of evolutionary young (diverged within the last 70 years) allopatric geothermal and ambient ecotypes. The four geothermal and ambient ecotype pairs were sampled to identify genomic patterns relating to thermal habitat divergence regardless of demographic history and age of divergence.

We sampled individuals across the four geothermal/ambient pairs for whole genome sequencing (n = 109). Each geothermal and ambient ecotype had roughly fourteen individuals (GTS geothermal = 14; GAR ambient = 14; MYV geothermal = 12; MYV ambient = 13; ASHN geothermal = 14; ASHN ambient = 14; SKR geothermal = 14; SKR ambient = 14). Genomic DNA was extracted from each individual from fin clips using the Quigene DNA extraction kit and were run on the Novoseq 6000 at the Center for Genomic Research at the University of Liverpool (SKR, GTS, GAR) and a HiSeqX at the University of Edinburgh (MYV, ASHN). Reads were mapped to the stickleback genome (Nath et al., 2021) using BWA-MEM(version 0.7.5a) (Li & Durbin, 2009) with the default parameters. Alignments were then filtered to remove reads with a mapping quality lower than 10, which equates to a 90% chance that the read was derived from another genomic location. Read duplicated were identified and filtered to retain a single representative using the Picard “MarkDuplicates” tool, version 1.85 (http://picard.sourceforge.net/). Variant detection was performed using the GATK HaplotypeCaller tool (McKenna et al., 2010; DePristo et al., 2011). Variants were then filtered using the GATK VariantFiltration tool (McKenna et al, 2010) following assessments of GATK recommended cut-offs. i.e. SNPS with either QD < 2, QUAL < 30, FS > 60, MQ < 40, SOR > 3, MQRankSum < -12.5 or ReadPosRankSum < -8 were flagged ”, and indels with QD < 2, QUAL < 30, FS >200, ReadPosRankSum < -20 were flagged. Single nucleotide polymorphisms (SNP) were filtered for minor allele frequencies less than 0.05 using vcftools (Danecek et al., 2011), and pruned for linkage disequilibrium using the –indep 50 5 2 flag in plink 1.9 (Purcell et al., 2007; Chang et al., 2015). After the filtering steps, we were left with 173,485 biallelic SNPs which were polymorphic in at least a single population.

### Population genetic structure based on whole genomes

We characterized fine scale population genetic structure and admixture across the four geothermal/ambient ecotype pairs. First, we determined the genetic relationship among all populations using a principal component analysis (PCA). The PCA was performed using pcadapt (Luu et al., 2017) which is based on Mahalanobis distance and a MAF cut-off of 0.01. Second, we assessed the amount of admixture and distinct genetic structure between the geothermal and ambient morphs from all populations using ADMIXTURE (Alexander et al., 2009). We tested K values from 1:10 with a cross validation procedure and the default parameters of ADMIXTURE. The optimal K value was determined as having the lowest cross validation error. Third, we calculated Fst between all geothermal and ambient ecotype pairs to estimate the degree of overall genomic differentiation of the thermal ecotypes using the stamppFST function from the StAMPP package (Pembleton et al., 2013). Lastly, we used Treemix (Pickrell & Pritchard, 2012) to establish the phylogenetic relationships among populations as well as the amount of introgression between populations. We ran Treemix for 1:10 migration edges and determined the greatest log likelihood of admixture events.

### Local adaptation to geothermal habitats

We identified regions of the genome that were potentially under selection and differentiated geothermal and ambient ecotypes. We calculated per locus Fst between geothermal and ambient ecotypes using plink (version 1.9) (Purcell et al., 2007; Chang et al., 2015). We then identified Fst outlier loci as any locus within the top 0.5% of the Fst distribution. In order to keep Fst outliers that are biologically meaningful we further filtered the outlier loci by only keeping loci that were greater than the average Fst of the outlier distribution for each geothermal/ambient ecotype pair. We then compared the degree of overlap of Fst outliers between geothermal/ambient ecotype pairs. We then identified the number of Fst outliers that corresponded to gene (coding) regions for each geothermal/ambient ecotype pair. Lastly, we assessed the degree of overlap in outlier genes between each ecotype pair and performed a gene ontology (GO) analysis for each ecotype pair using Metascape (Zhou et al., 2019).

### Parallel adaptation to geothermal habitats

We identified divergent regions of the genome which were common to all geothermal/ambient ecotype pairs. To minimize the amount of noise across the genome we performed a sliding window analysis using 25 Kilobase (Kb) windows with an overlap between windows of 1Kb using the R package windowscanr (github.com/tavareshugo/WindowScanR). We then filtered the dataset to include windows containing at least three SNPs within the window and defined Fst outliers as being in the top 0.5% of the Fst distribution and have a Fst value greater than the average Fst from the outlier distribution.

We then identified regions of the genome diverging in parallel between all geothermal and ambient adapted ecotypes. We conducted a scan to determine parallel changes in allele frequencies across all geothermal and ambient adapted ecotypes using an allele frequency vector analysis of parallel evolutionary responses, implemented using the R package AF-vapeR (Whiting et al., 2022). Each chromosome was split into 50 SNP windows and a matrix of allele frequencies were calculated for each window. Each cell within the matrix represented a change in allele frequency between geothermal and ambient ecotypes for a given SNP. An allele frequency vector was calculated for all SNPs within a given window and then the allele frequency vector was normalised so that all population pairs had equivalent variance.

Correlation coefficients were then calculated between the normalised allele frequency vectors and eigenvector decomposition was performed on the correlation coefficients for a given 50 SNP window. The changes in allele frequency vectors were then compared to a null distribution made up of randomized allele frequency changes across the populations. Full parallelism of allele frequencies were indicated as elevated values on the first eigenevector while multi-parallelism and divergent allele frequencies were indicated as elevated values on eigenvectors two and three.

Any chromosome showing significant parallel or multi-parallel changes in allele frequencies among geothermal and ambient adapted morphs were assessed for selective sweeps. We assessed regions of the genome that appeared to be diverging in parallel (or multi-parallel) trajectories for selective sweeps using the R package rehh (Gautier & Vitalis, 2012). We used the rehh package as it allows for cross population comparisons of extended homozygosity haplotypes based on the commonality of allele frequencies. We used the scan_hh function to calculate major and minor allele frequencies and the integrated haplotype homozygosity score (iHH) for each marker along a chromosome. As we did not know which markers were derived or ancestral we specified that the markers were not polarized allowing for the use of allele frequencies. We then used the function ies2xpehh to calculate extended homozygosity haplotypes between geothermal and ambient morphs based on allele frequencies, a Bonferroni correction was applied. Generally, values above 0 were associated with geothermal ecotypes and values below 0 were associated with ambient ecotypes.

Regions of the genome which showed parallel or multi-parallel changes in allele frequencies were also investigated to be associated with putative structural variants such as inversions. Local principal component analyses (PCA) have been shown to accurately identify structural variants across the genome (Li & Ralph, 2019; Mérot et al., 2020). We used the lostruct R package (Li & Ralph, 2019) to identify outliers in population structure along chromosomes containing significant and (multi) parallel changes in allele frequencies. Local PCA methods (i.e. sliding window analyses) have been shown to reliably detect structural variants, such as inversions, from short read sequencing data (Li & Ralph, 2019; Mérot et al., 2020). We used a window size of 50 SNPs and used the eigen_windows function to perform the local PCA along the chromosome containing the region of interest. We then used the pc_dist function to calculate a distance matrix between all windows in the previous analysis. Multidimensional scaling analysis, using the cmdscale function, was applied to the distance matrix to identify which PCA windows were population structure outliers.

Lastly, we identified all genes within regions that showed parallel or multi-parallel changes in allele frequencies and performed a gene ontology analysis. Gene ontology analysis can be used to identify terms that associated with the input genes as well as biological processes. We used Metascape (Zhou et al., 2019) to identify significant gene ontology terms associated with our genes of interest.

### Appetite, mass, and fat content phenotypes

To assess phenotypic variation relevant to our genomic findings we conducted appetite feeding trials, food restriction experiments, and assessments of body fat in fish from both geothermal and ambient habitats. For appetite trials we used lab bred F2 generation fish from ASHN, MYV, and SKR. We reared fish to a mature stage and following a fast of 24 hours we fed individuals (n = 18 per ecotype per lake) standard aquaculture pellets (EWOS, Weybridge, Surrey, Start 040 inc Boosterfeed, pellet size 0.9mm) to satiation. Satiation was defined as the point at which fish stopped feeding or were unable to consume more food. We counted the number of pellets that each individual ate as well as the time take to eat to the point of satiation. We used a type II ANOVA, using the aov function in the stats package, to test for the effects of population, ecotype, and their interaction on the number of pellets eaten to satiation as well as the time to satiation (model 1: Number of pellets eaten ∼ Population * Ecotype; model 2: Time to satiation ∼ Population * Ecotype).

Our tests for differences in weight loss under a restricted diet were conducted on wild fish from ASHN and MYV ecotype pairs. We collected between 17-26 individuals per ecotype per lake (41 ambient and 50 geothermal individuals total) in the summer of 2017. We then measured the initial total mass of each fish and tagged them with visible implant elastomers to enable individual identification before subjecting them to a reduced food intake for 30 days.

After the 30 day period we weighed each individual to calculate the amount of weight lost by each individual. We tested for differences in the amount of mass lost between geothermal and ambient ecotypes due to the treatment with a repeated measures ANOVA using the anova_test function in the rstatix package (https://github.com/kassambara/rstatix).

To test for differences in body lipid content between wild ecotypes caught in summer and winter of 2017 from five populations in Iceland. We collected between 18-20 individuals per ecotype in the summer and between 20-32 individuals per ecotype in the winter from ASHN, SKR, GTS, GAR, and STN (a location in the north of Iceland with geothermal stickleback). Collected stickleback were euthanized by phenoxyethanol overdose and fixed in 10% formalin in the field. We used diethyl ether to extract fat from each individual to assess fat content across different seasons (Hewavitharana et al., 2020). Individuals were dehydrated and weighed before being submerged in diethyl ether within a 50ml falcon tube for a period of 7 days to dissolve fat. Following this, individuals were once again dehydrated and weighted to obtain a difference in weight. We then used an ANOVA to model variation in lipid content due to population, ecotype, and season (Lipid content ∼ Population * Ecotype * Season) using the aov function in the stats package.

