## Supplementary for "Parallel adaptation to geothermally-warmed habitats due to common structural variation and functional developmental pathways"

### **Supplementary tables and Figures**

**Table1.** Sites sampled for threespined stickleback for the phylogeographic analysis.

| **Site name** | **Site code** | **Coordinates** |
| --- | --- | --- |
| Grettislaug | GTS | 65.88224 -19.73661 |
| Reykir | RKRW | 65.47086 -19.358078 |
| Causeway | CSWY | 65.73074 -19.462626 |
| Holmavatn | HGRNS | 65.675167 -19.486183 |
| Saudarkrokur Cold | SKRC | 65.732191 -19.618574 |
| Saudarkrokur Warm | SKRW | 65.732260 -19.618937 |
| Steinsstadir | STNST | 65.469154 -19.358097 |
| Myvatn Warm | MYVTNW | 65.633991 -16.923241 |
| Myvatn Cold | MYVTNC | 65.630196 -16.991621 |
| Kleifarvatn Cold | KLFC | 63.93811 -21.94195 |
| Thingvallavatn Warm | THNGW | 64.15059 -21.200521 |
| Litlaa | LITA | 66.076947 -16.684749 |
| Litlaa Pond | LITP | 66.082295 -16.681183 |
| Skjalftavatn | SKAL | 66.077446 -16.676031 |
| Reykholt Warm | RKLTW | 64.6618 -21.374348 |
| Reykholt Cold | RKLTC | 64.67892 -21.2506 |
| Ashildarholtsvatn Warm | ASHNW | 65.72516 -19.600725 |
| Ashildarholtsvatn Cold | ASHNC | 65.724976 -19.601784 |
| Grimsstadir | GRMT | Unknown (collected by Bjarni) |
| Reykir River Cold Site | RKRC | Unknown |
| Behind barn on farm | BARN | 65.4764 -19.3697 |
| Ljosaland | NH | 65.4761 -19.3748 |
| Thingvallavatn Cold | THNGC | Unknown |
| Opnur Warmer end | OPNURW | Unknown |
| Opnur Colder End | OPNURC | Unknown |

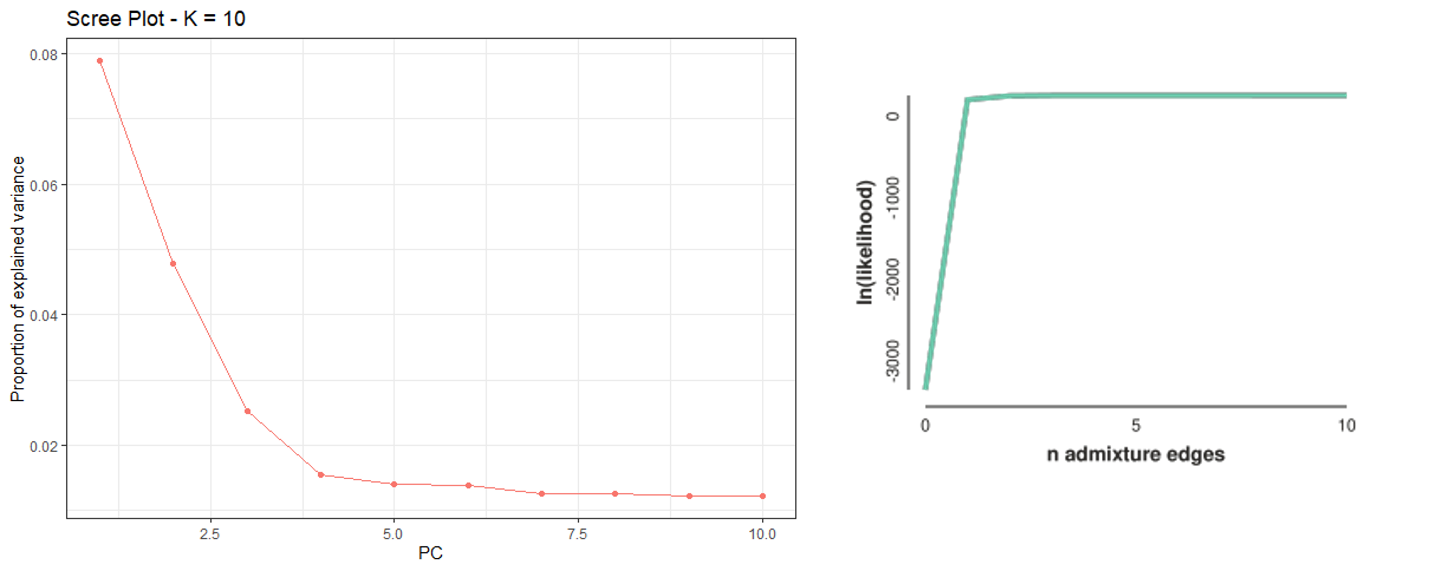

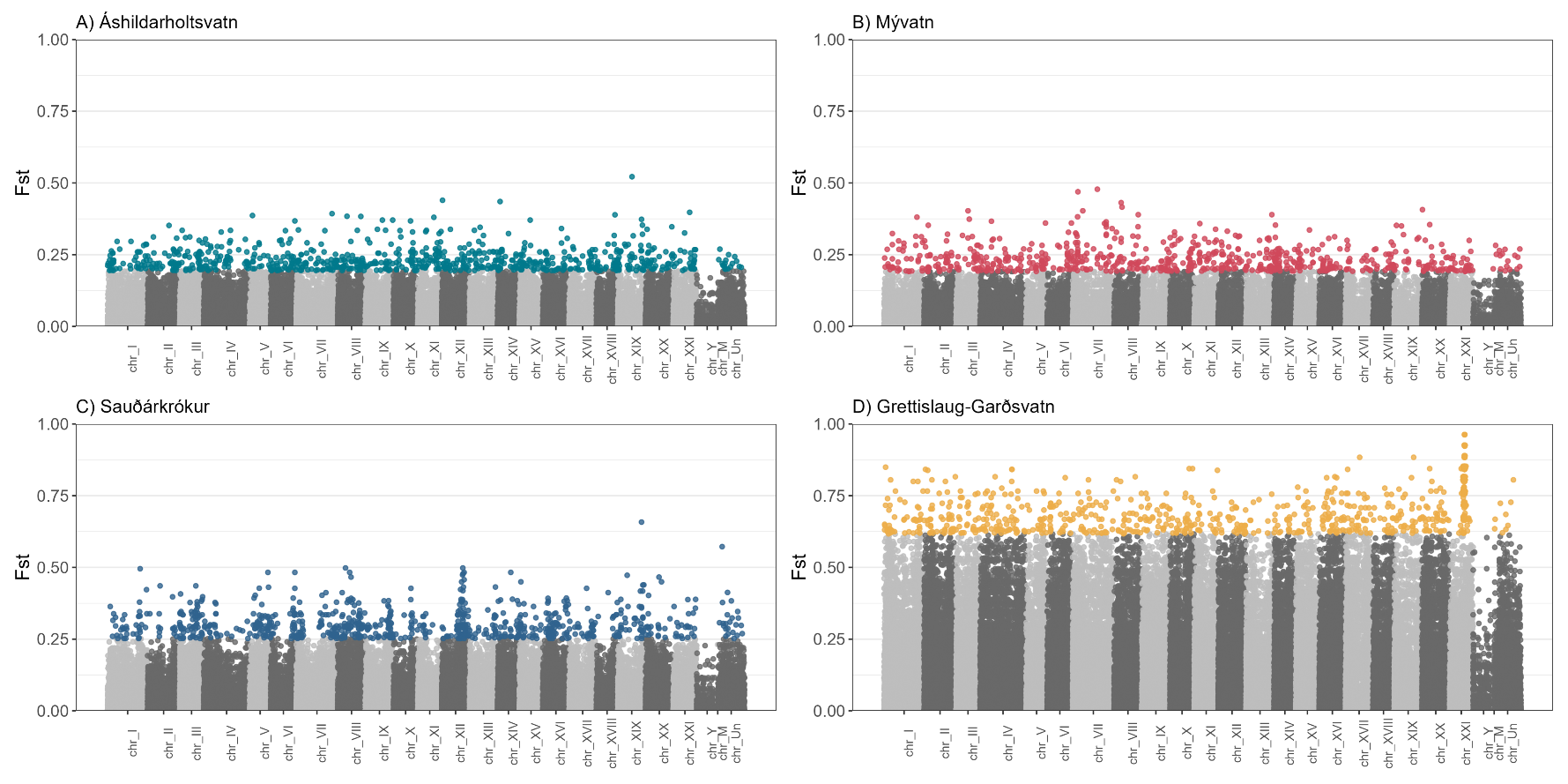

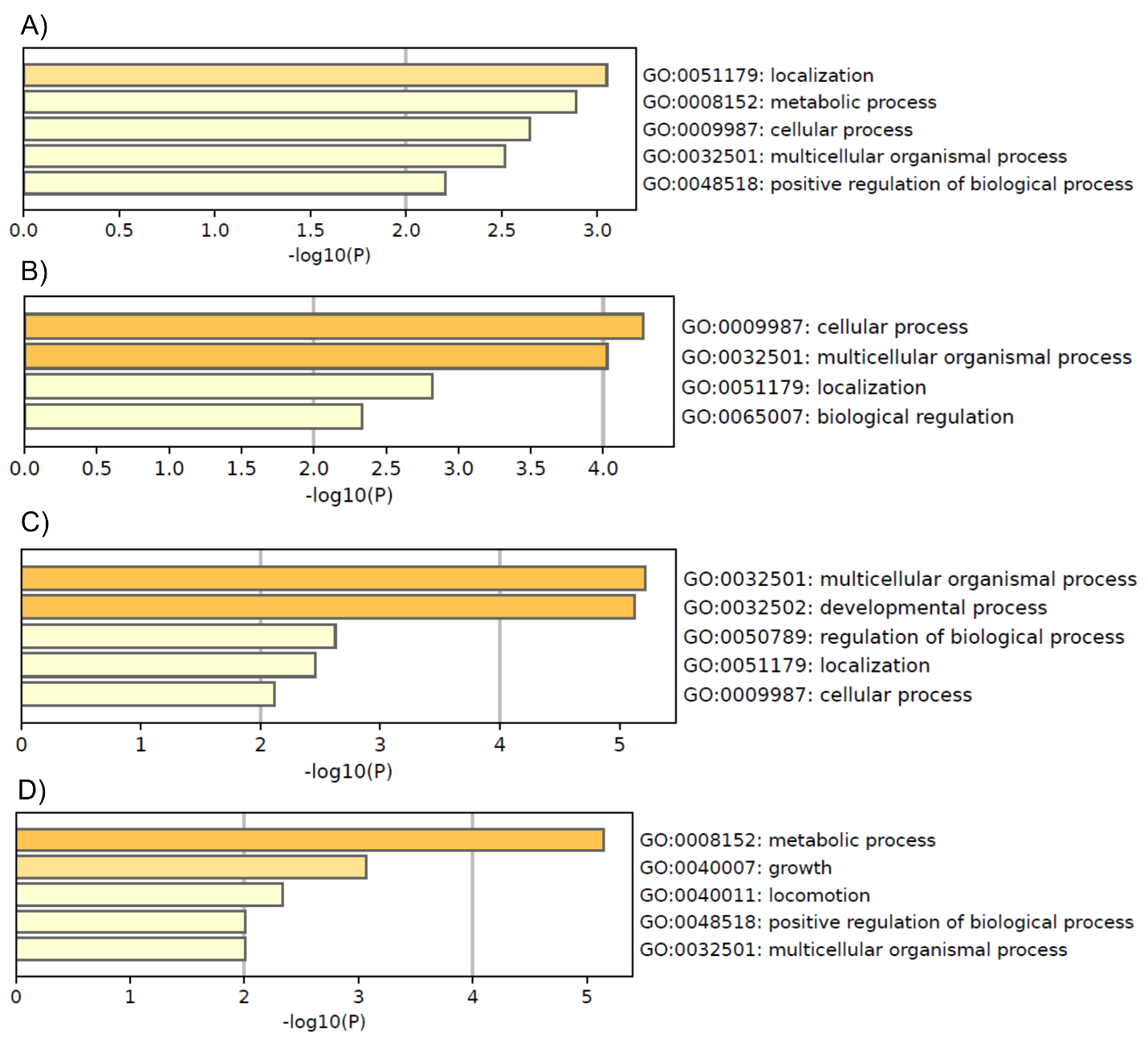

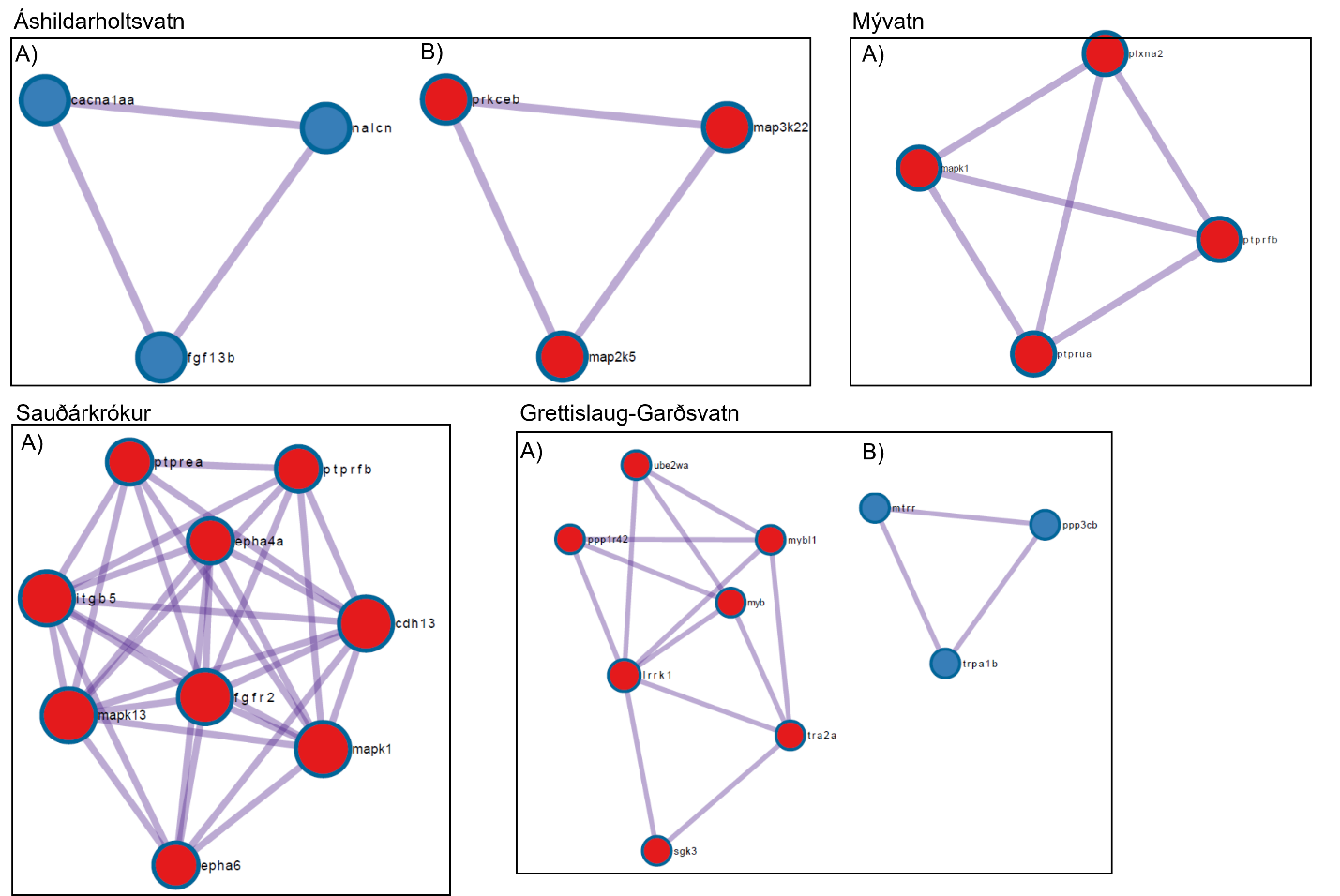

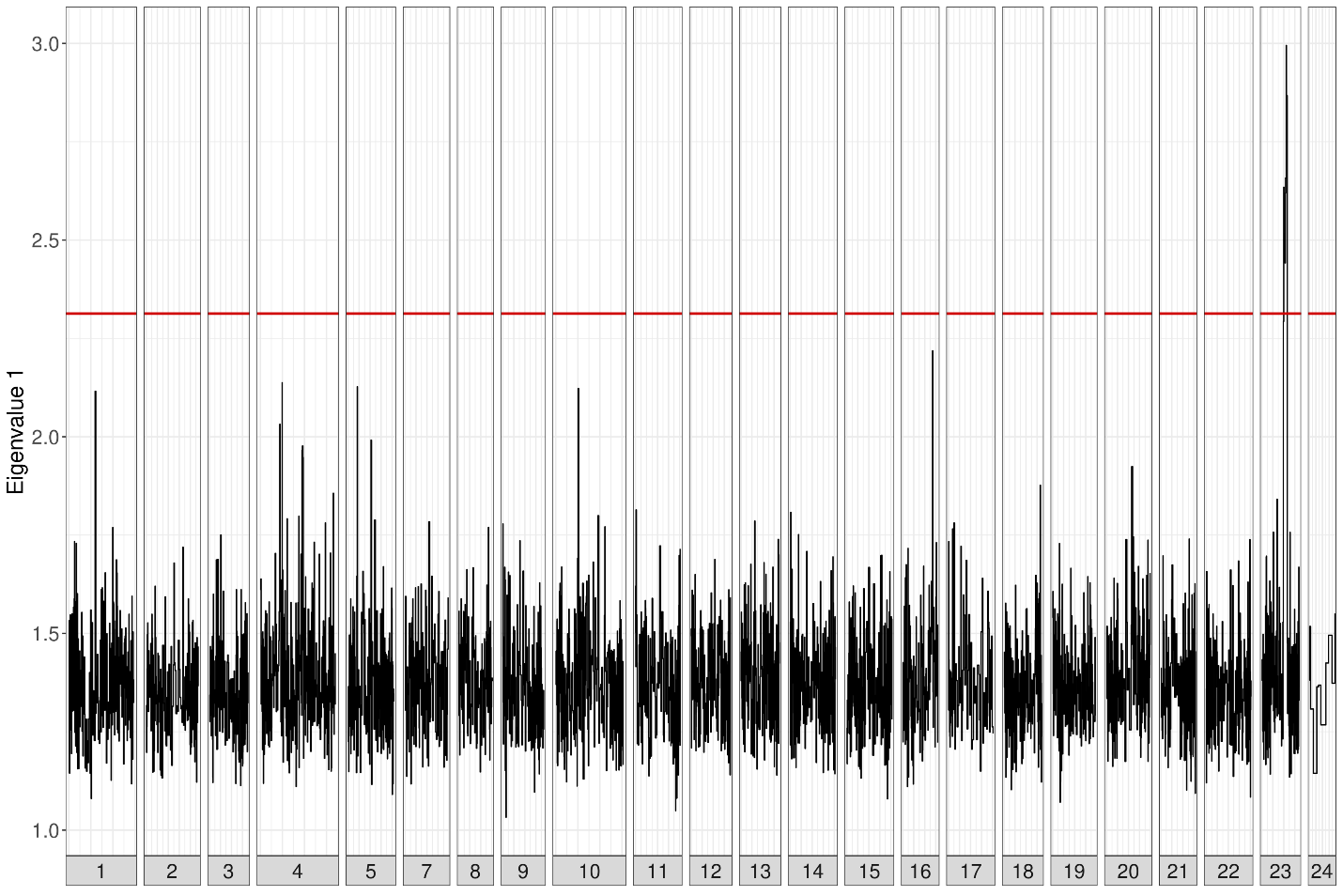

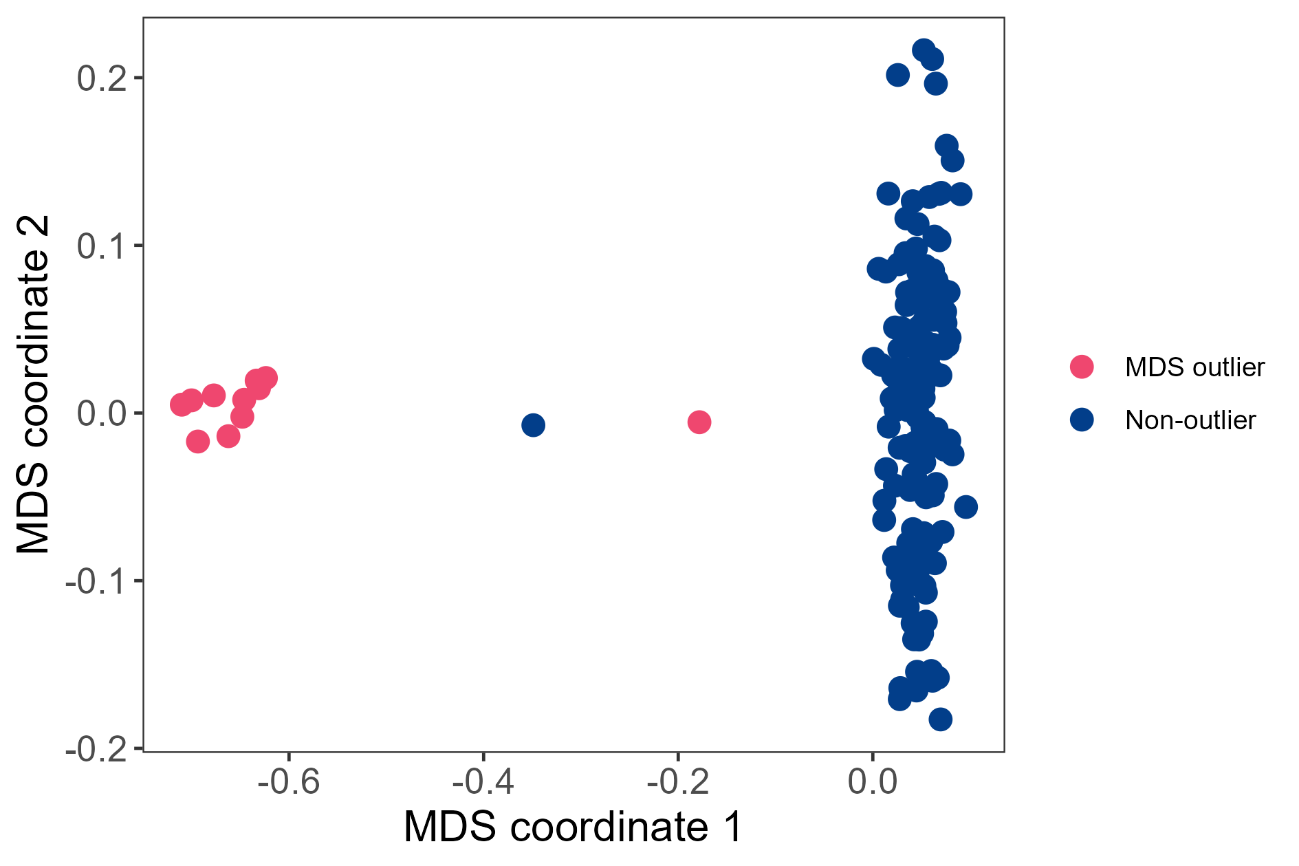

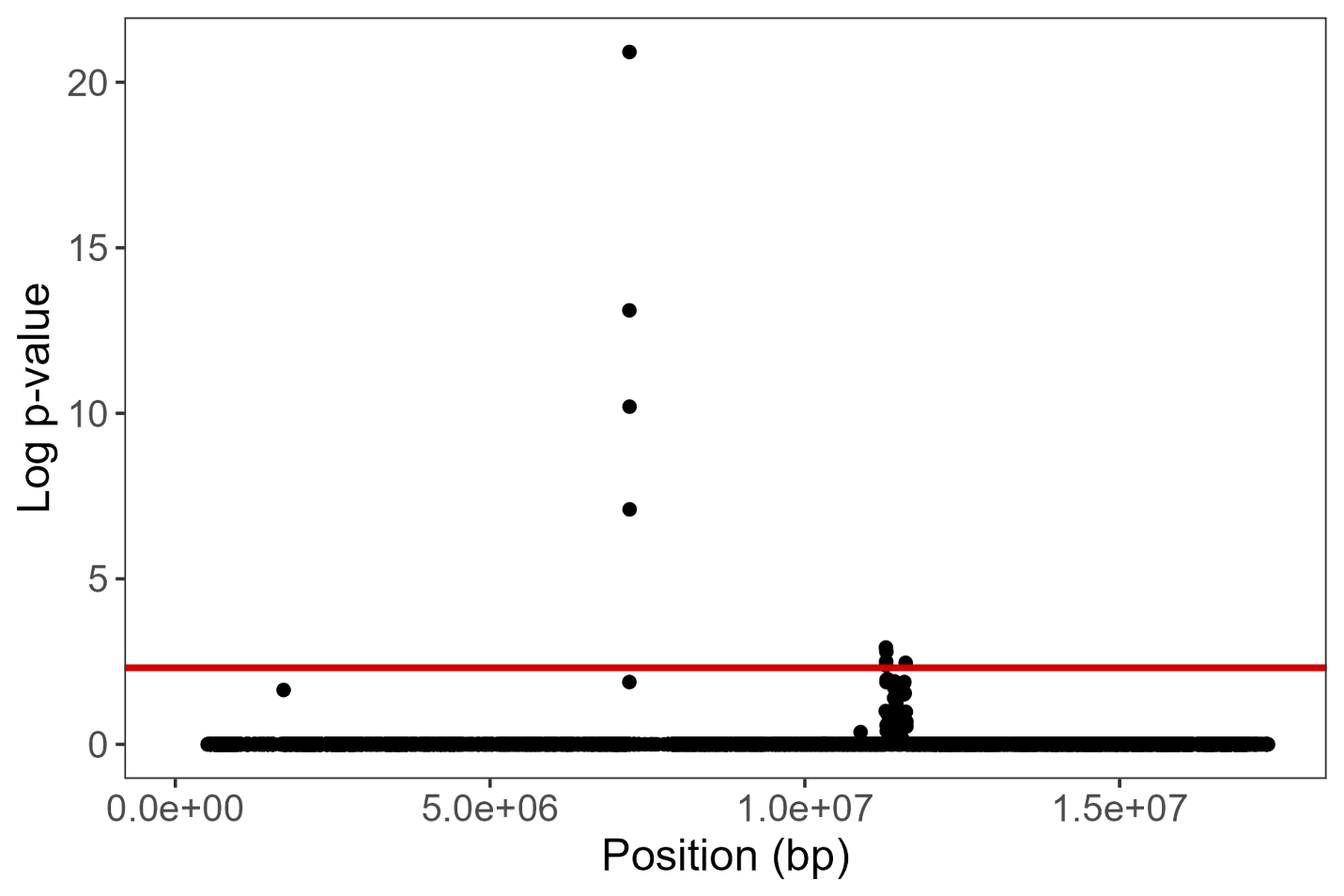

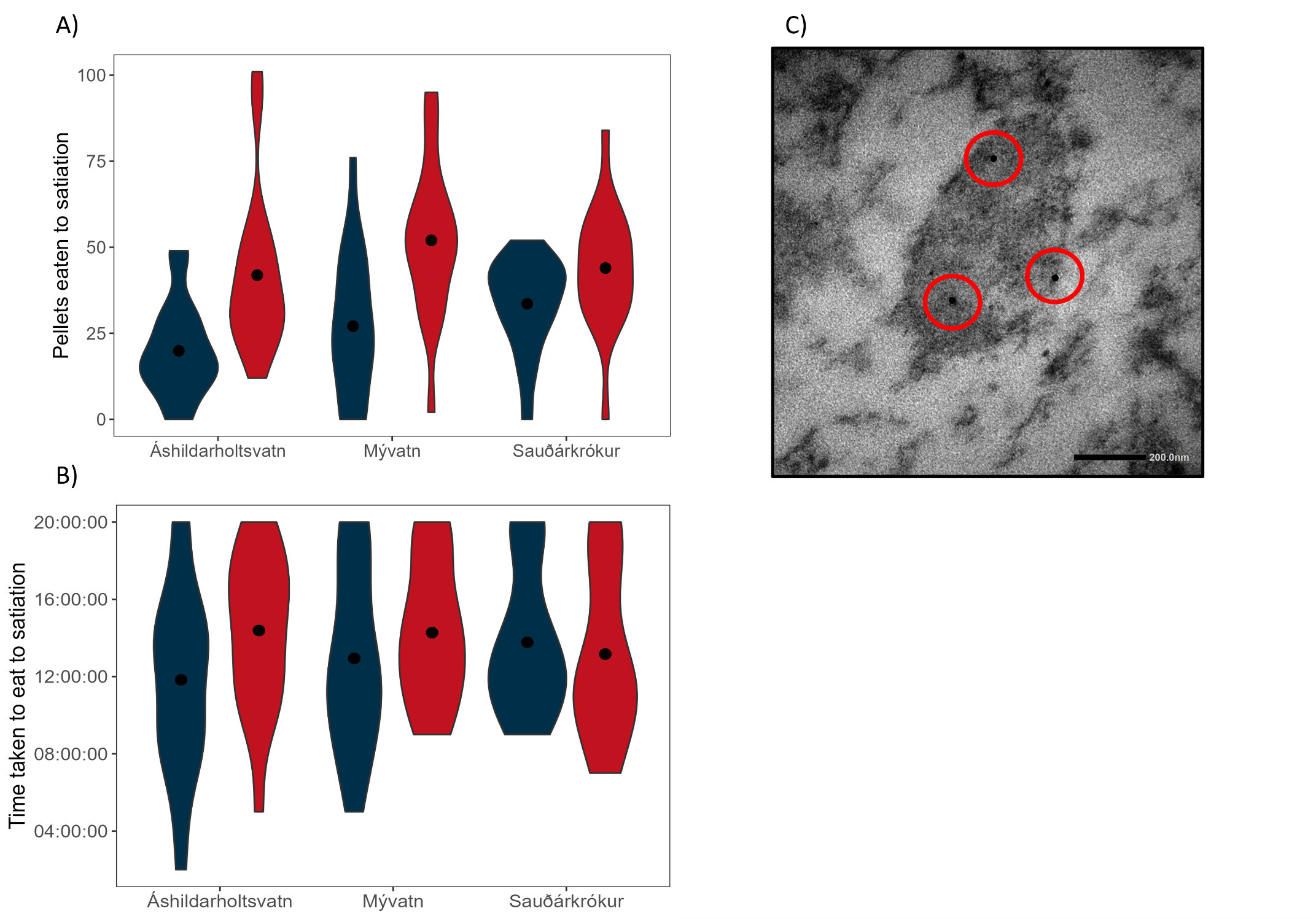
